# Hippocampal Dipeptidyl Peptidase 9 Bidirectionally Regulates Memory Via Synaptic Plasticity

**DOI:** 10.1101/2023.11.02.565088

**Authors:** Ya-Bo Zhao, Shi-Zhe Wang, Wen-Ting Guo, Le Wang, Xun Tang, Jin-Nan Li, Lin Xu, Qi-Xin Zhou

## Abstract

It has been reported that peripherally expressed subtypes of the dipeptidyl peptidase (DPP) family, such as DPP4, modulate memory. However, interestingly whether DPP9 which one of the central nervous systems (CNS) enriched isoforms, regulates memory has not been elucidated yet. Here, we report that DPP9, which is found almost exclusively in neurons, is highly expressed and has high enzyme activity in many brain regions, especially in the hippocampus. Hippocampal DPP9 expression increases after fear memory formation. Fear memory was weakened by DPP9 knockdown and enhanced by DPP9 protein overexpression in the hippocampus. According to subsequent hippocampal proteomics, multiple pathways were enriched by DPP9 expression changes, including the peptidase pathway, which can be bidirectionally regulated by DPP9, and pathways involved in the regulation of synaptic plasticity. DPP9 interacts with its enzymatic substrate neuropeptide Y (NPY) in neurons directly. Hippocampal long-term potentiation (LTP), a form of synaptic plasticity, further confirmed the key role of DPP9 in decreasing LTP through DPP9 knockdown and enhancing LTP through its overexpression. Moreover, inhibiting DPP9 enzyme activity impairs both plasticity and memory. Besides, Affinity purification mass spectrometry (AP-MS) revealed that DPP9-interacting proteins are involved in the functions of dendritic spines and axons. By combining AP-MS and proteomics, DPP9 was shown to play a role in regulating actin functions. Taken together, our findings reveal that DPP9 affects the CNS not only through enzymatic activity but also through protein‒protein interactions. This study provides new insights into the molecular mechanisms of memory and DPP family functions.

## Introduction

The dipeptidyl peptidase (DPP) family contains four prolyl-specific peptidases: DPP4, fibroblast activation protein (FAP), DPP8 and DPP9[1]. DPP4 reportedly ameliorates AD-related memory loss, although DPP4 and FAP are expressed mainly peripherally [2], which suggests that the DPP family can indirectly affect the central nervous system (CNS) [3]. However, the functions of the isoforms, such as DPP9, which are mainly expressed in the CNS, have not been determined.

Allen brain data show that DPP9 mRNA is widely distributed in the CNS [4]. DPP9 has post-proline dipeptidyl aminopeptidase activity that cleaves Xaa-Pro dipeptides from the N-termini of proteins [5]. It plays a biological role by cleaving specific substrates, such as neuropeptide Y (NPY), peptide YY (PYY), and glucagon-like peptide-1 (GLP1) [6]. NPY is a neurotransmitter widely distributed in the brain. NPY can affect learning and memory by interacting with NPY receptors and initiating signaling cascades[7].

The hippocampus is a vital brain region located in the medial temporal lobe and is responsible for learning and memory. It plays a crucial role in the formation and retrieval of declarative memories, such as facts, events, and personal experiences [8]. Synaptic plasticity is widely recognized as a cellular mechanism of memory in which synapses can strengthen or weaken in response to neural activity [9]. Long-term potentiation (LTP) is a form of synaptic plasticity. LTP initiates structural changes in synapses through the cytoskeletal protein actin [10]. Actin dynamics contribute to the formation and remodeling of synaptic connections, thereby affecting synaptic plasticity[11]. However, the molecular mechanisms underlying memory are not yet fully understood [12, 13].

In this study, we found that DPP9 is highly expressed almost exclusively within neurons in the hippocampus and that its expression in the dorsal hippocampus bidirectionally regulates fear memory in mice. This study demonstrated that DPP9 regulates several synaptic and memory-related proteins and that its enzymatic activity affects memory. Moreover, we found that DPP9 interacts with NPY in neurons and that the actin regulatory protein can regulate synaptic plasticity. These findings suggest that DPP9 plays a critical role in fear memory.

## Materials and Methods

### Animals

Adult male C57BL/6J mice (from SPF (Beijing) Biotechnology Co., Ltd., 6-8 weeks of age, weighing 18–22 g) were used. At the Kunming Institute of Zoology, animals were housed in cages with four animals each and provided with free access to food and water under a 12-hour light/dark cycle in a temperature-controlled environment. Tunnel handling was performed every day for 4 days before the behavioral experiment, and the tunnel was used when the mice were transferred in the behavioral experiment[14]. The experimental procedures were approved by the Animal Ethics Committee of the Kunming Institute of Zoology (No. IACUC-RE-2021-06-012).

### Stereotaxic injection

Mice were anesthetized with Zoletil 50 (40 mg/kg) and xylazine (10 mg/kg) and fixed in a stereotaxic instrument (RWD, China). The soft tissue was removed from the skull after the scalp was cut. The skull was microdrilled after the skull was sutured. The coordinates of the hippocampus were 1.7 mm along the AP axis, ±1.5 mm on the medial-dorsal (ML) axis, and 1.8 mm in dorsal-ventral (DV) depth with respect to bregma. A nanoliter-volume injection pump (Micro4, WPI, USA) was used to inject 500 nl of virus (at an injection speed of 50 nl/min) into each hippocampus. After the injection, the glass pipette was left for 5 minutes and removed. Finally, the mouse’s scalp was sutured, and the mice were returned to their home cages. Ibuprofen (10 mg/kg) was given for pain relief within 3 days after the operation.

### DPP9 knockdown and overexpression

Knockdown and overexpression were performed using AVV vectors (Shandong Vigene Biosciences Co., Ltd.). The DPP9 KD group was injected with AAV2/9-CMV-Cax13d-FLAG-U6-2x gRNA (DPP9) mixed with AAV2/9-hSyn-GFP-WPRA. The gRNA sequences used for DPP9 were 3’CCCCATAGACAAAGAGCACAGTG5’ and 3’AGTCTCGATGTTTGCCCACCCACA5’; the DPP9 KD control group was injected with AAV2/9-CMV-Cax13d-FLAG-U6-2x gRNA (randomly) mixed with AAV2/9-hSyn-GFP-WPRA. The DPP9 OE group was injected with AAV2/9-hSyn-DPP9-FLAG mixed with AAV2/9-hSyn-GFP-WPRA; the DPP9 OE control group was only injected with AAV2/9-hSyn-GFP-WPRA.

### Immunofluorescence staining

Mice were perfused with 50 ml of phosphate-buffered saline (PBS) after anesthesia, followed by 150 ml of 4% PFA. The brains were kept in 4% PFA at 4°C overnight and dehydrated twice using 30% sucrose. The brains were frozen after being embedded in OCT compound. For immunofluorescence staining, 15 μm slices were washed 3 times with PBS containing 0.3% Triton X-100 (PBST). The slices were placed in citrate buffer solution (Beyotime, P0081) for antigen retrieval (microwave oven, low power, 20 min). Sections were blocked with 1.5% donkey and goat serum (Solarbio, SL050, SL038) in PBST (blocking buffer) after they had returned to room temperature for 1.5 h. Slides were incubated with primary antibodies (rabbit anti-DPP9, Abcam, ab42080, 1:250; mouse anti-NeuN, Millipore ABN78B, 1:500; mouse anti-NPY, Santa Cruz, sc-133080, 1:500) diluted in blocking buffer for 24 h at 4°C. After incubation, the cells were washed with PBST 3 times. The secondary antibodies (donkey anti-rabbit Alexa Fluor Plus 488, Invitrogen, A32790, 1:1000; goat anti-mouse Alexa Fluor Plus 555, Invitrogen, A32727, 1:1000) were diluted with blocking buffer, and the slices were incubated at room temperature for 1 h. After washing 3 times with PBST, the slides were incubated with DAPI (Beyotime, C1005) and then mounted with ProLong Glass Antifade Mountant (Invitrogen, P36980). After the mounting medium was allowed to dry for 10 h, the cells were photographed with an FV3000 laser confocal microscope (Evident, Japan).

### Western blotting

The mouse brain was quickly removed 21 days after virus injection, and hippocampal tissues were placed in ice-cold PBS. The tissue was homogenized in RIPA buffer supplemented with protease inhibitors (Thermo, 87785) for 1 min on ice and then centrifuged at 10,000 × g for 10 min at 4°C. The supernatants of the samples were assayed for protein content using a Pierce Rapid Gold BCA kit (Thermo, A53225), diluted with LDS sample buffer (Invitrogen, NP0007) and 50 mM DTT and then heated at 70°C for 10 min. The proteins were loaded onto 10% PAGE gels and transferred onto PVDF membranes. The membranes were blocked with QuickBlock Blocking Buffer (Beyotime, P0220). The membranes were incubated with the following primary antibodies overnight at 4°C: mouse anti-DPP9 (Origene, TA503937, 1:1000). After three washes with TBST, the membranes were incubated with anti-mouse H+L HRP-conjugated IgG (Proteintech, SA00001-1, 1:10000) for 1 h at room temperature, washed with PBS, and scanned with a chemiluminescence detection system (Tanon, China).

### DPP8/9 enzyme activity

A DPPIV-Glo Protease Assay Kit (Promega, G8350) was used for DPP9 enzyme activity analysis. Briefly, the hippocampus was homogenized in TE buffer (pH 8.0) supplemented with 10 μM sitagliptin and centrifuged at 10,000 × g for 10 min at 4°C[15]. The supernatant was diluted 50 times. The substrate and buffer were mixed to prepare the reaction buffer. The diluted supernatant and reaction buffer were mixed in a 96-well plate and incubated at 37°C for 20 min. A microplate reader (Allsheng, China) was used to measure the chemiluminescence intensity.

### Proximity Ligation Assay (PLA)

Following the protocol of the Duolink In Situ Detection Reagents Orange (Sigma, DUO92007), PLA experiments were also conducted. Briefly, sections were blocked with donkey and goat serum after permeabilization and antigen retrieval. Primary antibodies (rabbit anti-DPP9, Abcam, ab42080, 1:250; mouse anti-NPY, Santa Cruz, sc-133080, 1:500) were added to the blocking buffer and incubated overnight at 4°C. After washing with PBST 3 times, anti-mouse (MINUS) and anti-rabbit (PLUS) probes were added and incubated for 2 h at room temperature. The samples were washed 3 times and incubated with ligation and ligation buffer at 37°C for 30 min. After washing, the polymerase was incubated with the amplified products at 37°C for 100 min. The slides were incubated with DAPI (Beyotime, C1005) and then mounted with ProLong Glass Antifade Mountant (Invitrogen, P36980). After the mounting medium was allowed to dry for 10 h, the cells were photographed with an FV3000 laser confocal microscope (Evident, Japan).

### Electrophysiological records

For LTP recordings of DPP9 KD and OE mice, brain slices (350 μm) were removed 21 days after the mice were injected with AAV. After the mice were anesthetized with isoflurane, the brains were quickly removed and placed in ice-cold saturated carbogen (95% O2, 5% CO2) artificial cerebrospinal fluid (ACSF) (120 NaCl, 2.5 KCl, 26 NaHCO3, 1.25 NaH2PO4, 2 MgSO4, 10 D-glucose and 2 CaCl2, mM). After slicing with a vibratome (Leica, German), the brain slices were placed in ACSF supplemented with carbogen at 30°C for 30 min and then maintained at room temperature for at least 1 hour. The brain slices were placed in the immersion chamber, and ACSF was continuously perfused with carbogen at a flow rate of 2-4 ml/min. The stimulating electrodes (made of a pair of wound Teflon-coated platinum-iridium alloys [WPIs]) were placed on Schaffer collaterals (SCs); the recording electrodes (made of borosilicate glass capillary drawn, 1 MΩ, PG10165, WPI) were placed in the stratum radiatum of the dorsal CA1 area of the hippocampus. The signals were recorded for at least 20 min at 0.033 Hz to establish a stable baseline with the stimulus intensity set to elicit an equivalent maximum fEPSP slope of approximately 40-50%. LTP was then induced by high-frequency stimulation (HFS, 3 columns of 1 s stimulation at 100 Hz with a 20 s interval between columns) at the same stimulus intensity as that used for baseline recordings. For recordings using inhibitors, brain slices were placed in ACSF supplemented with carbogen-containing inhibitors (Val-BoroPro, MedChemExpress, HY-13233A; Sitagliptin, MedChemExpress, HY-13749B) 30 min before placement in the chamber, and the ACSF perfused in the chamber also contained inhibitors. Signals were acquired using a Multiclamp 700B and Digidata 1440A (Molecular Devices, USA) and digitized at a sampling rate of 20 kHz.

### Novel object recognition test

This experiment was performed as previously described[16]. Briefly, the day before the NOR test, the mice were placed in an empty box (50 × 50 × 40 cm, white PMMA) to allow them to move freely for 10 minutes, during which they were allowed to habituate to the context. Two identical objects (Lego blocks) were placed on both sides of the box 5 cm away from the wall, and the mice were allowed to move freely through the box for 10 minutes. After 24 hours, the mouse was returned to the box after the old object from the previous day was replaced with a new object (plaster cast). The time for the mice to explore the new object and the old object was recorded separately. After each mouse experiment, the box was thoroughly cleaned with 75% ethanol. The preference index was calculated by subtracting the exploration time of the old object from the exploration time of the new object and dividing by the total exploration time (the exploration time of the new object plus the exploration time of the old object).

### Open field test

The mice were placed in an open field device (Med Associates, Inc., MED-OFAS-MSU) and allowed to move freely for 15 minutes. After each mouse experiment, the box was thoroughly cleaned with 75% ethanol. The data were analyzed using Activity Monitor 7 software.

### Elevated Plus Maze

The mice were placed in the experimental room for at least 1 hour before the experiment. The mice were placed on the Elevated Plus Maze (Med Associates, Inc., ENV-560A) and allowed to move freely for 5 min. After each mouse experiment, the box was thoroughly cleaned with 75% ethanol. A stopwatch was used to record the time the mice spent in the open and closed arms.

### Contextual fear conditioning

The day before the plantar electric shock, the mice were placed in the Fear Conditioning System (Med Associates, Inc., MED-VFC2-USB-M) for 10 minutes to allow the mice to habituate to the context. The next day, the mice were placed in the fear conditioning system. For the DPP9 OE mice, we administered 3 foot shocks (0.5 mA, two minutes apart); for the DPP9 KD mice, we administered 5 foot shocks (0.8 mA, session of shock stimulus interval (s): 150, 90, 80, 100, 70). After the shock, the mice were returned to their home cages. After 48 hours, the mice were returned to the fear conditioning system for 5 minutes to allow memory retrieval. VBP intervention in the conditioned fear memory experiment The cannula was embedded in the hippocampus of the mice 7 days before adaptation, and 1 μL of 500 μM VBP was given 10 minutes before 5 kHz sound training. Give 1 μL of 500 μM VBP 10 min before 5 kHz memory retrieval.

### Cannula placement

Cannula placement is similar to stereotaxic injection. After the mouse was anesthetized, it was placed in the stereotaxic instrument. The skull was microdrilled after the skull was sutured. The coordinates of the hippocampus were 1.7 mm along the AP axis, ±1.5 mm on the medial-dorsal (ML) axis, and 1.6 mm in dorsal-ventral (DV) depth with respect to bregma. The cannula was placed at the target site and fixed with denture base resin. The wound was disinfected using iodophor. The mice were returned to their home cages. Ibuprofen (10 mg/kg) was given for pain relief within 3 days after the operation.

### Proteome sample preparation and LC‒MS

Sample preparation was performed using filter-aided sample preparation (FASP)[17]. Briefly, the hippocampal tissue of the mice was removed 4 hours after the CFC experiment. The tissue was homogenized in lysis buffer (100 mM NaCl, 50 mM Tris, 0.5% Triton X-100, 0.5% SDS, pH 7.4) containing protease inhibitors (Thermo, 87785) for 1 min on ice and then centrifuged at 10,000 × g for 10 min at 4°C. The supernatants of the samples were assayed for protein content using a Pierce Rapid Gold BCA kit (Thermo, A53225). Fifty micrograms of protein was added to DTT (Aladdin, D104860) to a final concentration of 20 mM and heated at 90°C for 10 min to reduce the protein concentration. Afterwards, iodoacetamide (Aladdin, I131590) at a final concentration of 40 mM was added for alkylation at 37°C in the dark for 30 min. Then, 5 times the volume of UA buffer (8 M urea, 100 mM Tris, pH 8.0) was added, and the mixture was placed into a 30K ultrafiltration tube (Pall Life Sciences, OD010C34) for cleaning (12000xg centrifugation for 15 min). The ultrafiltration tube was cleaned with UA buffer 3 times. Then, 0.5 μg of Lys-C (Wako, 125-05061) was added, and the mixture was incubated at 37°C for 4 hours. Then, 1 μg of trypsin (Promega, V5280) was added, and the mixture was digested at 37°C for 8 hours. The samples were desalted with a custom-made stage tip (ion exchange SDB-RPS, Empore, 2241) after enzymatic digestion[18]. The samples were dried using a Speedvac and reconstituted with 0.1% formic acid in water. The concentration of the peptide was measured using the absorbance at 280 nm. The peptide solution (800 ng) was injected into an UltiMate 3000 RSLC nanosystem (Thermo, USA) with a C18 25 cm × 75 μm column (Thermo, 164941). The mass spectrometer used was a Q Exactive (Thermo, USA). The peptides were eluted with 2–28% buffer B (0.1% [vol/vol] formic acid, 80% acetonitrile) with a nonlinear 90 min gradient at a flow rate of 300 nL/min.

The samples were analyzed on a Q Exactive mass spectrometer with the Data Independent Acquisition (DIA) method for peptide MS/MS analysis. Full scans of peptide precursors were performed from 350 to 1490 m/z at 70 K FWHM resolution with a 3 × 106 AGC target and a maximum IT of 100 ms. After a full scan, a variable window was used to acquire DIA from 150 m/z at 17.5 K FWHM resolution (at 200 m/z) with a 1 × 106 AGC target and a maximum IT of 100 ms. HCD fragmentation was applied with 26% collision energy. All the data were searched using a spectral library free of DIA-NN 1.8.1, and the precursor FDR was set at 1.0%[19].

### Affinity purification mass spectrometry (AP-MS)

The hippocampus was removed 21 days after the mice were injected with the AVV of DPP9 OE. The tissue was homogenized in lysis buffer (100 mM NaCl, 50 mM Tris, 0.5% NP-40, 10% glycerin, pH 7.4) containing protease inhibitors (Thermo, 87785) for 1 min on ice and then centrifuged at 10,000 × g for 10 min at 4°C. Mouse anti-FLAG antibody (Sigma, F3165) and Protein G magnetic beads (10004D, Invitrogen) were added to the supernatant, which was subsequently incubated at 4°C for 3 hours. The magnetic beads were washed 4 times with PBST buffer. The 3XFLAG peptide (MedChemExpress, HY-P0319A) was added for competitive elution. The eluate was added to LDS sample buffer (Invitrogen, NP0007) and 50 mM DTT and then heated at 70°C for 10 min. The proteins were loaded onto NUPAGE gels (Invitrogen, NP0322), and the gels were stained with silver stain (Beyotime, P0017S) after electrophoresis. The gel slices were excised for in-gel digestion[20]. Briefly, gel slices were reduced and alkylated with DTT and iodoacetamide after destaining. After adding 1 μg of trypsin overnight at 37°C, acetonitrile was added to extract the peptides. A homemade STAGE tip was used for desalting. The samples were dried using a Speedvac and reconstituted with 0.1% formic acid in water. The peptide solution was injected into an EASY-nLC 1000 (Thermo, USA) with a custom-made C18 20 cm × 75 μm column. Then, 2–28% buffer B (0.1% formic acid, 100% acetonitrile) was used to elute the peptides with a nonlinear 60 min gradient at a flow rate of 200 nl/min.

The samples were run on an Orbitrap Fusion Lumos (Thermo, USA) mass spectrometer with the Data Dependent Acquisition (DDA) method for peptide MS/MS analysis. Full scans of peptide precursors were performed from 350 to 1600 m/z at 60 K FWHM resolution with a 4 × 105 AGC target and a maximum interval (IT) of 50 ms. After a full scan, MS2 signals from 100 m/z at 15 K FWHM resolution with a 5 × 104 AGC target and a maximum interval (IT) of 22 ms were applied. HCD fragmentation was applied with 30% collision energy. The original data files were retrieved and analyzed with Proteome Discoverer 2.4 software, and the FDR was set to 0.1%. The mass spectrometry proteomics data have been deposited in the ProteomeXchange Consortium via the iProX partner repository[21, 22] with the dataset identifier PXD044715.

### Statistical analyses

The statistical data were analyzed with GraphPad Prism (version 9.2.0) and R (version 4.2.3). The illustration was drawn with www.biorender.com. Numerical data are presented as the mean ± SEM. Statistical significance was analyzed using Student’s t test for two groups and analysis of variance (ANOVA) for three or more groups. Two-way ANOVA was used to analyze differences in the learning curves of the behavioral paradigms. For all analyses, p < 0.05 was considered to indicate statistical significance.

## Results

### DPP9 is widely distributed in neurons

It can be seen in public databases that DPP9 mRNA is widely distributed in the brain, but it is unclear whether the protein distribution is consistent or if the cellular localization is consistent[23]. To investigate the distribution of the DPP9 protein in the brain, we performed immunofluorescence staining for DPP9 and NeuN, a neuronal marker, in sections of the mouse brain (Fig 1. a). DPP9 was highly expressed in the hippocampus (HIP), cerebellum (CBM), cortex (COX), and olfactory bulb (OB) (Fig 1.b and Supplemental Fig 1.c-e). We measured DPP9 enzymatic activity in different brain regions. The highest enzymatic activity was found in the hippocampus, and olfactory bulb had the activity in the thalamus and hypothalamus was relatively low (Figure 1c). DPP9 positive (DPP9+) cells were highly colocalized with NeuN-positive (NeuN+) cells in brain areas with high DPP9 enzymatic activity. In the hippocampus and cortex, DPP9 is expressed only in neurons, while in other areas, such as the cerebellum and olfactory bulb, DPP9 is expressed in non-neuronal cells. (Figure 1d, n=3; F (3, 16) =7.238; HIP vs. CBM, *** p<0.001; HIP vs. OB, *** p<0.001; two-way ANOVA). DPP9 also displayed some non-cell-specific signals in the slices, suggesting that it is secreted outside the cells. As a post-proline cleaving serine protease enzyme, it is a key indicator of DPP9 function [24]. These findings suggested that DPP9 may be involved in neuronal activities.

**Figure 1.**
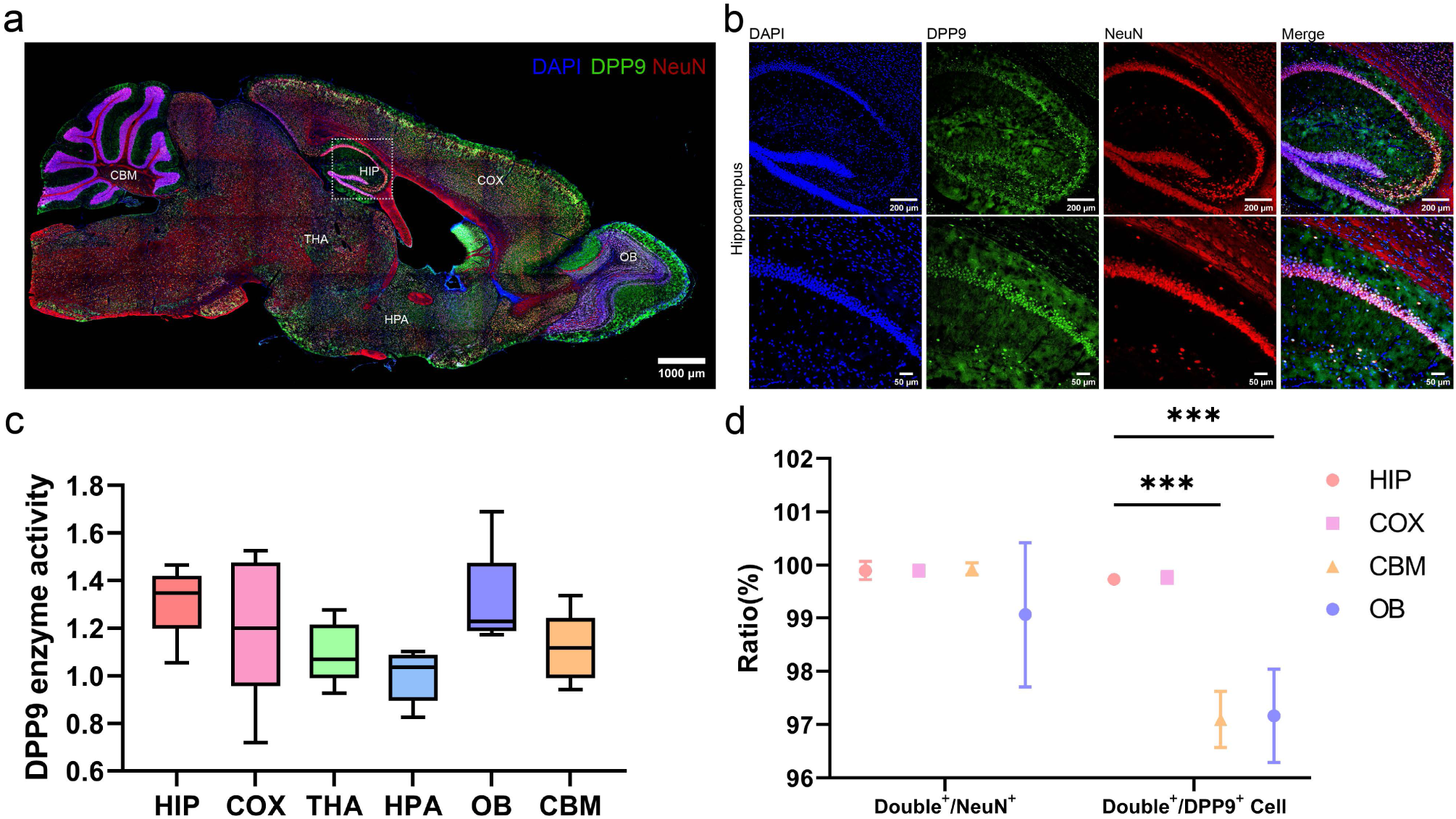
Expression pattern of DPP9 in the mouse brain and expression of DPP9 after memory retrieval. (a) Immunofluorescence staining of mouse brain tissue. Sagittal sections (ML: 1.00 mm) for DPP9 (green), NeuN (red), and DAPI (blue) staining; scale, 1000 μm. (b). Immunofluorescence staining of the mouse hippocampus. DPP9 (green), NeuN (red), DAPI (blue), 10X (upper) scale, 200 μm; 20X (lower) scale, 50 μm. (c). Enzyme activity of DPP9 in different brain regions: the cortex (COX), hippocampus (HIP), thalamus (THA), hypothalamus (HPA), olfactory bulb (OB), and cerebellum (CBM) (n=5; one-way ANOVA) (d). The DPP9+ and NeuN+ merged cell (double+) and DPP9+ cell ratios in the hippocampus, cerebellum, cortex, and olfactory bulb (n=3, t test). *** P<0.001, mean ± SEM.

### Fear memory increases hippocampal DPP9 expression

We then detected DPP9 protein levels 48 h after context fear conditional memory retrieval. The expression level of DPP9 was significantly increased (Fig 2. b, n=5; **p=0.042; t test), suggesting that DPP9 may play an important role in learning and memory. A CasRx-based DPP9 knockdown (KD) vector and a DPP9 overexpression (OE) vector were designed to generate two adeno-associated viruses (AAVs) (Fig 2. c, f). We first verified the efficiency of the knockdown sgRNAs in cells (Supplemental Fig. 2 a-d). Subsequently, we injected DPP9-KD AAV along with GFP AAV into the dorsal hippocampus of mice to ensure the accuracy of the injection position. After 21 days of expression, perfusion samples were collected for immunofluorescence staining, and western blot analysis was performed to confirm the knockdown efficiency (Fig 2. d). The results showed a significant decrease in DPP9 expression in the dorsal hippocampus (Fig 2. e, n=3, *p=0.010, t test). Similarly, we injected DPP9 OE AAV into the dorsal hippocampus of mice and observed a significant increase in DPP9 expression after 21 days (Fig 2. h, n=3; **p=0.002; t test) (western blot raw file in Supplement Fig. 6). Since exogenously expressed DPP9 contains a FLAG tag, we used FLAG staining to distinguish endogenous DPP9. Therefore, we successfully established DPP9 knockdown and overexpression models using AAV-mediated gene delivery.

**Figure 2.**
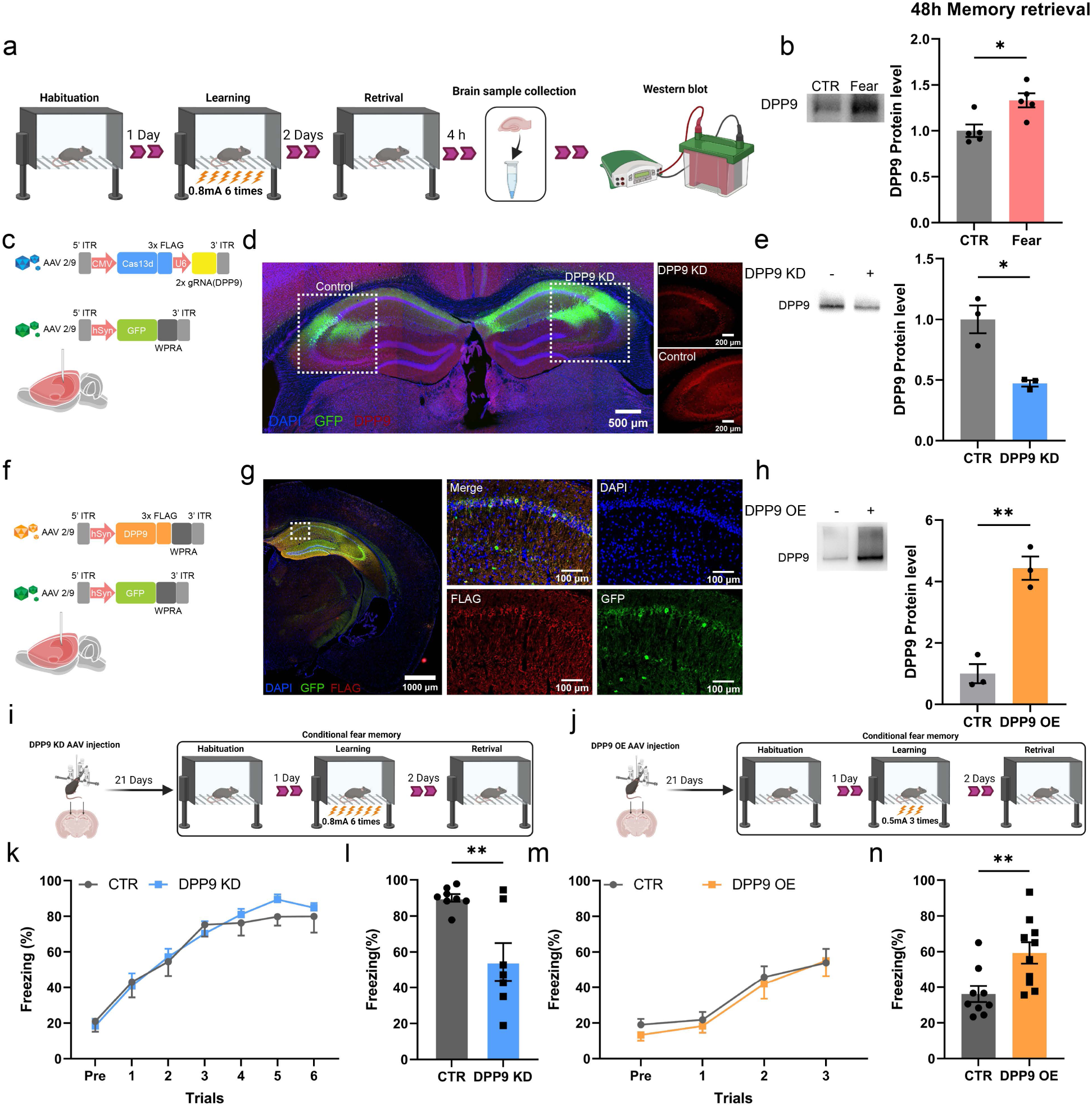
The DPP is regulated by memory and regulates memory (a). Schematic of contextual fear conditioning (CFC) and DPP9 protein detection after memory retrieval. (b). Immunoblotting for DPP9 in the hippocampus 48 h after foot shock. (c) Schematic of the AAV vector delivering Cas13d and gRNA. (d) Immunofluorescence staining of injection sites 24 h after KD AVV injection (GFP, green; DPP9, red; DAPI, blue); scale, 1 mm; 100 mm. (e) Immunoblotting for DPP9 in the hippocampus 24 h after KD-AVV injection (n=3, t test). (f) Schematic of the AAV vector used to deliver DPP9-FLAG. (g) Immunofluorescence staining of injection sites 24 h after OE-AVV injection (GFP, green; FLAG, red; DAPI, blue); scale, 1 mm; 100 mm. (h) Immunoblotting for DPP9 expression in the hippocampus 24 h after OE-induced AVV injection (n=3, t test). (i, j) Schematic of the CFC after KD and OE AAV injection. (k) Learning curve during CFC training (6 times, 0.8 mA) for KD mice. (n = 8, RM two-way ANOVA) (m) Learning curve during CFC training (3 times, 0.5 mA) for OE mice. (n = 10, two-way ANOVA). (l, n) Memory retrieval of CFCs on day 2 in KD and CTR mice (n = 7, t test). *P<0.05, ** P<0.01, mean ± SEM.

### DPP9 expression bidirectionally regulates long-term memory maintenance but not the acquisition process

Since the hippocampus is critical for learning and memory and DPP9 is highly expressed in the hippocampus, we investigated the role of DPP9 in these processes. To reduce animal use, we conducted a series of behavioral tests on the same mice. Tests were separated by at least 24 hours to minimize interference, and tests were started on those that had weaker stimulation. We used open-field experiments to examine whether DPP9 affects spontaneous behavior and locomotion in mice. We found that DPP9 KD and OE mice did not exhibit changes in movement speed or distance (Supplemental Fig 3. a, n=12, p=0.892; b, n=12, p=0.780; c, n=8, p=0.183; d, n=8, p=0.839; t test). We then conducted novel object recognition experiments to test long-term memory and found that DPP9 KD mice exhibited significant impairment in their ability to distinguish new objects from old objects after 24 hours (Supplemental Fig 3. g, n=11, **p=0.003, t test), while DPP9 OE mice sniffed new objects significantly longer than did the control group (Supplemental Fig 3. h, n=9, **p=0.003, t test). Next, we conducted an elevated plus maze test to detect anxiety in mice, and the results showed that DPP9 OE and DPP9 KD did not affect the time mice spent in the open arm (Supplemental Fig 3. e, n=12, p=0.263; f, n=11, p=0.971; t test). Finally, we used a conditioned fear test, specifically the contextual fear conditioning test, to simultaneously examine learning and long-term memory in mice. Since memory is enhanced after DPP9 OE, we designed different foot shock models for DPP9 OE and KD patients. We used weaker foot shocks (0.5 mA, 3 times) to detect memory enhancement after DPP9 OE (Fig 2.i) and stronger foot shocks (0.8 mA, 6 times) to detect memory impairment after DPP9 KD (Fig 2.j). Notably, neither KD nor OE of DPP9 affected the learning performance of the mice (Fig 2. k, n=8; F (1, 14) =0.079, p=0.781; m, n=10; F (1, 19) =0.286, p=0.598; two-way ANOVA). The results of memory retrieval 48 hours after learning were consistent with those of the novel object recognition experiment, and the freezing behavior of DPP9 KD mice was significantly reduced (Fig 5, l, n=7, **p=0.003; n, n=9, **p=0.007; t test); moreover, the freezing behavior of DPP9 OE mice was significantly increased. In conclusion, our results showed that knocking down or overexpressing DPP9 in the dorsal hippocampus does not affect spontaneous movement, anxiety or fear learning but strongly regulates fear memory retrieval.

### Proteomic analysis revealed that DPP9 regulates memory-related pathways

We demonstrated that DPP9 plays a significant role in memory, but the mechanism by which DPP9 regulates memory is still unclear. We investigated the role of DPP9 in memory and identified DPP9-derived differentially expressed proteins (DEPs) via proteomics. Both the OE and KD and their controls experienced a strong learning model (CFC, 0.8 mA, 6 times), and 48 hours later, they experienced memory retrieval. Four hours later, bilateral hippocampal tissues were removed for enzymatic digestion, data-independent acquisition (DIA) and DIA-NN analysis (Fig 3. a). Principal component analysis (PCA) revealed that the principal components associated with DPP9 knockdown (KD) and overexpression (OE) partially overlapped with their respective controls, but most of them were separated, indicating that changes in the expression of DPP9 affect many proteins in the hippocampus (Fig 3. b, c). Subsequent volcano plot analysis also confirmed the PCA results. We found 539 DEPs with a P value <0.05 in the DPP9 knockdown protein group (Fig3. d, including 57 DEPs with a log (fold change) <0.5 and a log (fold change)>0.5 were included; the DPP9-overexpressing protein group contained 1354 DEPs with a P value < 0.05 (Fig 3. e), 240 DEPs with a log (fold change) <0.5 and 456 DEPs with a log (fold change)>0.5). Moreover, we used the top 100 proteins with the smallest P values to construct a hierarchical clustering heatmap and perform KEGG and GO pathway enrichment analyses (supplement Fig 4 a-e). Next, we used a Venn diagram to identify the DEPs in both the DPP9-KD and DPP9-OE samples and found 163 DEPs common to both groups (Fig. 3 f). We performed KEGG pathway and GO enrichment analyses on these DEPs and found that they were highly enriched in neural and synapse-related pathways (Fig. 3 g, h). Notably, according to GSEA, immune-related pathways were enriched, indicating that DPP9 may also regulate proteins related to neuroimmune functions (supplement Fig 4, f, g). Our findings provide a theoretical basis for how DPP9 affects memory by regulating proteins involved in neuronal and synaptic functions.

**Figure 3.**
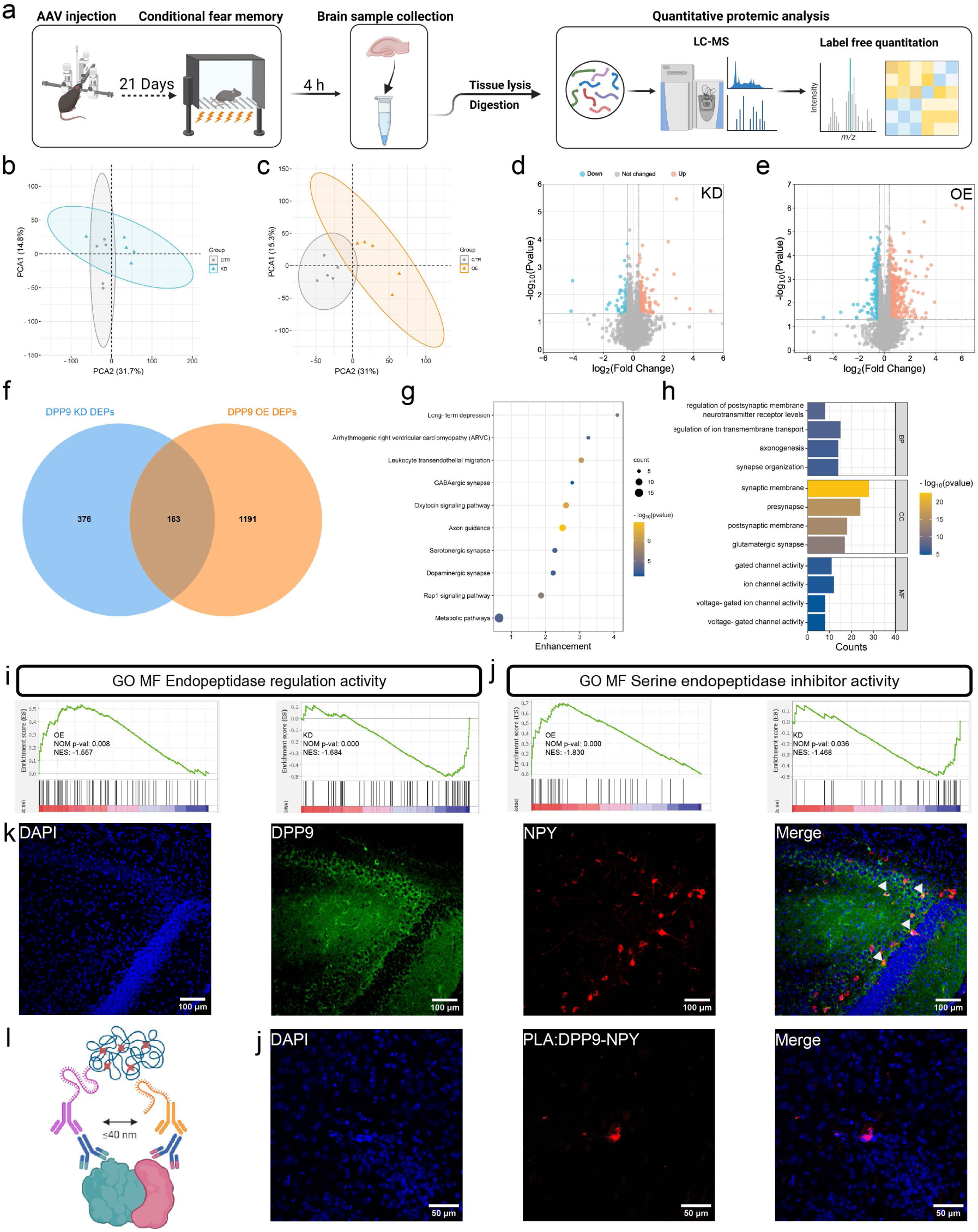
DPP9 proteomic characterization and regulation of NPY. (a) Schematic of the proteomic analysis after CFC retrieval in KD and OE mice. (b, c) Principal component analysis (PCA) plot of all proteins from KD and OE mice. (n=5) (d, e) Volcano plots showing the -Log10 P value versus fold change between KD or OE mice and CTRs. (f) Venn diagram of DPP9 KD DEPs and DPP9 OE DEPs; there are 163 common proteins. (g) KEGG pathway enrichment analysis of KD-OE comment DEPs. (h) GO enrichment analysis of KD-OE comment DEPs. (i) GSEA was used to construct a plot for GO:MF endopeptidase regulation activity; OE vs CTR (left); and KD vs CTR (right). (j) GSEA was used to construct a plot for GO enrichment: MF serine endopeptidase inhibitor activity, OE vs CTR (left), and KD vs CTR (right). (k) Immunofluorescence staining and colocalization of DPP9 and NPY in the hippocampus. DPP9 (green), NPY (red), and DAPI (blue) are shown. Scale=100 μm. (l) Schematic of the proximity ligation assay (PLA). (j) PLA for DPP9 and NPY in the hippocampus; PLA (red), DAPI (blue); scale=50 μm.

### DPP9 regulates memory through enzymatic activity and interacts with NPY

Then, we used GSEA to identify the signaling pathways in which the DPP9-OE and KD samples exhibited opposite trends. The endopeptidase regulation activity pathway was enriched (Fig 3.i), and we also enriched serine endopeptidase inhibitor activity (Fig 3.j), which indicated that DPP9 is a serine protease in the hippocampus. Neuropeptide Y (NPY), a substrate of DPP9, plays a key role in learning and memory. We therefore designed a series of experiments to prove the relationship between NPY and DPP9 in the hippocampus[7]. We found that DPP9 was co-expressed in NPY+ neurons in the hippocampus (Fig. 3. k). However, these findings do not prove that DPP9 interacts with NPY in the hippocampus because NPY is the substrate of DPP9, and detecting this interaction via traditional protein interaction methods (such as co-IP) is infeasible. We used a proximity ligation assay (PLA) to detect the interaction between DPP9 and NPY in situ in the hippocampus, and the results showed that direct protein-protein interaction occurred in the hippocampus (Fig. 3 i, j).

To further explore the effect of DPP9 and its enzymatic activity on memory, we first tested hippocampal LTP in mice that were injected with DPP9 KD or OE or control AAV for 21 days. Using three trains of 100 Hz high-frequency stimulation (HFS) to induce LTP in hippocampal slices, we found that LTP was significantly attenuated after DPP9 KD compared to that in KD control mice (Fig 4 a, b, n=7, **p=0.003; t test). In contrast, LTP was significantly enhanced after DPP9-OE mice than in OE control mice (Fig. 4 c, d; n=7; ***p<0.001; t test). After that, we utilized Val-BoroPro (VBP), a nonselective inhibitor of postproline cleaving serine proteases that inhibit DPP4, DPP8, and DPP9, to test the enzymatic activity of DPP9 on LTP. [25]. LTP was significantly weakened by 10 μM VBP (Fig 5g, h; n=6; *p=0.024; t test) and completely blocked by 20 μM VBP (Fig 4. e, f; n=6; **p=0.002; t test). To determine whether VBP is the cause of LTP damage through DPP4 inhibition, we used the DPP4-specific inhibitor sitagliptin to exclude the role of DPP4. We observed no change in HFS-induced LTP after high doses (50 μM) of sitagliptin were administered (Fig 4. i, j, n=6, p=0.684; t test).

**Figure 4.**
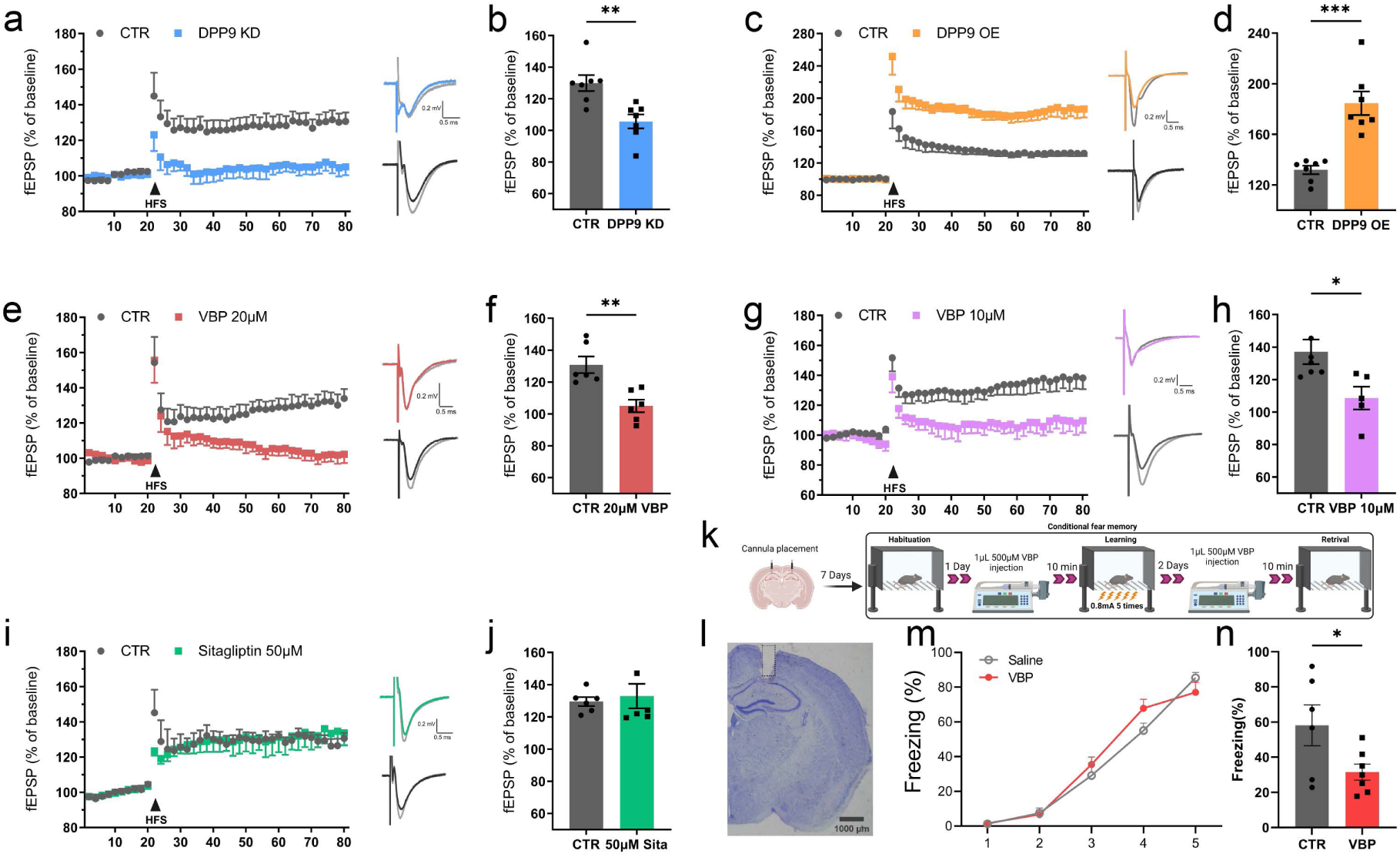
Effects of DPP9 and DPP9 inhibitors on long-term potentiation. (a-j) High-frequency stimulation (HFS, 100 Hz) induced CA1 long-term potentiation (LTP). The solution was maintained for at least 1 hour and incubated for 20 min for the fEPSP summary. (KD, n=7; OE, n=7; Val-BoroPro (VBP), 20 μM; n=6; VBP, 10 μM; n=6; sitagliptin, 50 μM; n=6; t test). (k) Schematic diagram of the VBP intervention in the conditioned fear memory test. The cannula was embedded in the hippocampus of the mice 7 days before adaptation, and 1 μL of 500 μM VBP was given 10 minutes before 5 kHz sound training. One microliter of 500 μM VBP was added 10 min before memory retrieval. (l) Nissl staining of brain sections at the cannula embedding site. (m) Learning curve during CFC training (4 times, 0.8 mA) for 500 μM VBP. (n=6, two-way ANOVA) (n) The CFC memory test was performed after 2 days of treatment with 500 μM VBP. (n=6, t test) *P<0.05, **P<0.01, *** P<0.001, mean ± SEM.

**Figure 5.**
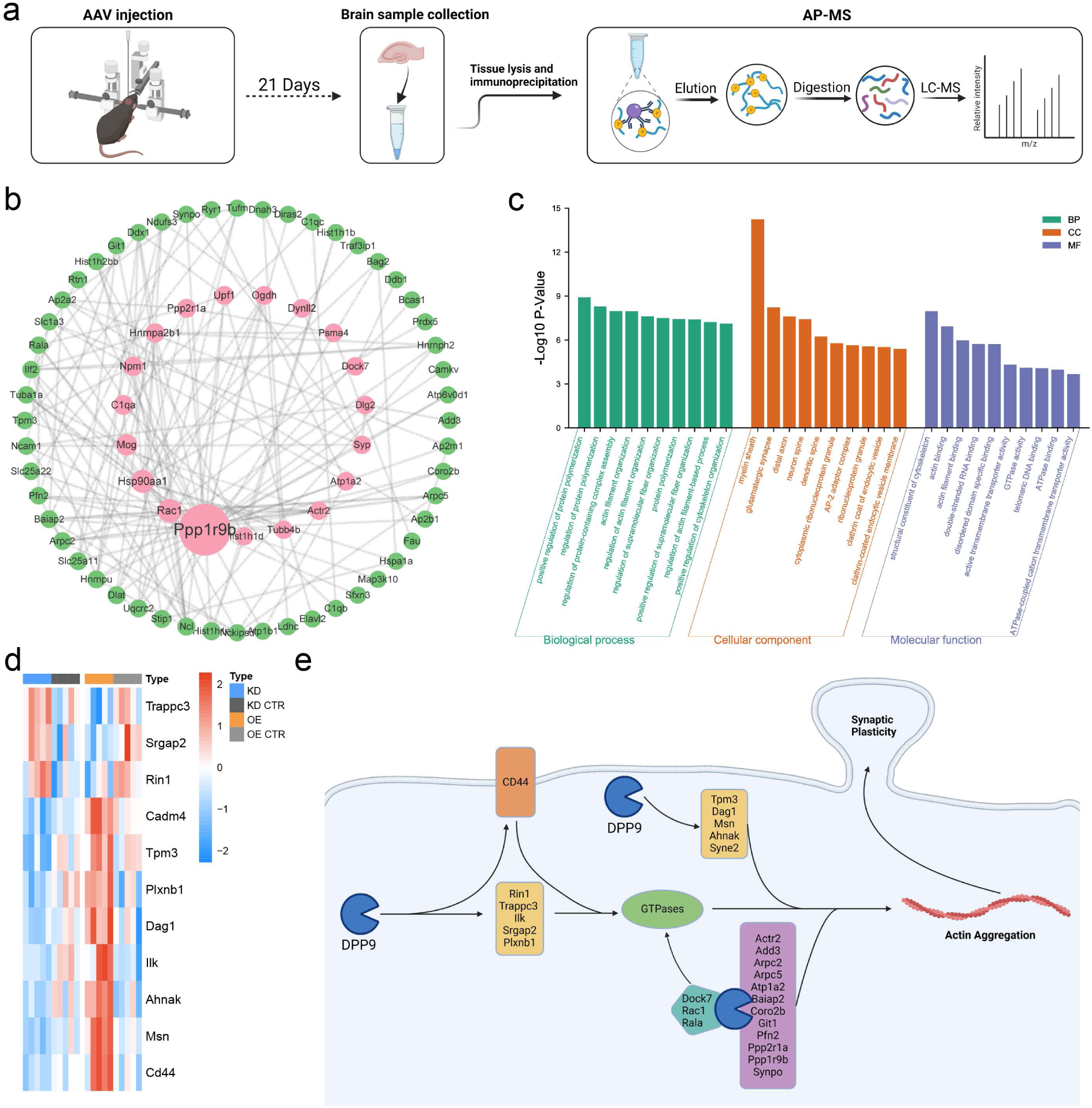
DPP9 regulates actin function. (a) Schematic of affinity purification and mass spectrometry (AP-MS) in the mouse brain. (b) STRING PPI analysis of DPP9-interacting proteins: pink (high interactions) and green (low interactions) (n=4). (c) DPP9 interaction protein-encoding gene ontology (GO) enrichment analysis. (d) Heatmap of KD-OE comment DEPs associated with small GTPases and actin. (e) Heatmap of KD-OE DEPs associated with the regulation and modification of postsynaptic function. (e) Schematic representation of how DPP9 indirectly regulates actin through a small GTPase and can also directly regulate actin.

Finally, we implanted the cannula in the hippocampus of the mice, performed memory retrieval 48 hours after the mice had experienced foot shocks, and injected 1 μL of 500 μM VBP 10 minutes before retrieval (Fig 4. k, i). We found that the freezing behavior of the mice was significantly reduced during memory retrieval (Fig 4. n, n=6; *P=0.031; t test) but not during the learning phase (Fig 4. m, n=6; F (1,11) =1.480; P=0.249; two-way ANOVA). The above experimental results proved that DPP9 is involved in the NPY process and that the enzyme activity and expression of DPP9 can affect the memory and synaptic plasticity of mice.

### DPP9-interacting protein and proteomic data revealed that DPP9 regulates actin function

Although the enzymatic activity of DPP9 is important, some studies have shown that the protein interaction of DPP9 also plays a regulatory role [26]. To investigate the potential binding of DPP9 to other proteins in the hippocampus and its impact on memory, we conducted affinity purification mass spectrometry (AP-MS) analyses after coimmunoprecipitation (Co-IP) to identify the proteins that interact with DPP9 directly in this region. Specifically, we expressed FLAG-tagged DPP9 (DPP9-FLAG) in the hippocampus, used magnetic beads with a FLAG antibody to pull down DPP9-FLAG, digested the protein, and identified its interacting partners via mass spectrometry (Fig. 5. a). We analyzed four samples and selected proteins that were detected in at least two samples simultaneously. In total, we identified 71 proteins that interact with DPP9. To gain further insight into the functions of these proteins, we performed a STRING protein network interaction (PPI) analysis and performed betweenness analysis and sorting. We placed proteins with greater numbers of interactions in the inner circle and marked them in pink, while those with fewer interactions were placed outside the circles and marked in green (Fig 5. b). We also conducted GO enrichment analysis of the 77 proteins and found that they were strongly associated with protein polymerization and actin functions (Fig 5.c). Notably, actin functions are closely linked to synaptic plasticity and memory [11]. The proteome data suggest that proteins that are bidirectionally regulated by DPP9 are also highly involved in the regulation of actin function and small GTPases. We generated heatmaps of these related proteins (Fig 5.d). Finally, we combined AP-MS and proteomics data and found that DPP9 affects synaptic plasticity by affecting actin function (Fig. 5. e). We conclude that DPP9 plays an important role in memory retrieval not only by cleaving NPY enzymatically but also through its regulation of actin.

## Discussion

In the present study, we found that high DPP9 expression is widely distributed in the brain, especially in the hippocampus. Moreover, we found that downregulating DPP9 expression in the dorsal hippocampus of mice specifically impaired memory retrieval and LTP, but upregulating DPP9 expression increased memory retrieval and LTP. We identified memory- and synaptic-related pathways regulated by DPP9. DPP9 participates in the interaction of NPY and regulates memory and LTP through enzymatic activity. Finally, we demonstrated that DPP9 may act on synaptic plasticity by affecting actin aggregation.

Using immunofluorescence staining for DPP9 and NeuN, we observed that DPP9 is widely expressed in the brain and is likely present in neurons. These results were confirmed by FISH analysis of the Allen Brain. Interestingly, DPP9 was also found to be expressed in nonneuronal cells in the cerebellum and olfactory bulb, suggesting its involvement in significant functions in these brain regions. Subsequently, we found that DPP9 functions as a peptidase in the hippocampus. Although NPY was proven to be the substrate of DPP9 in previous studies, it was not clear whether DPP9 was involved in the regulation of NPY in vivo[27, 28]. Our study demonstrated that DPP9 directly interacts with NPY in the hippocampus and that inhibition of this enzyme can impair LTP and fear memory in mice. NPY is cleaved from full-length NPY 1-36 to NPY 3-36 by DPP9, and these two NPYs play different roles in the nervous system. NPY 1-36 mainly bind to the Y1, Y2 and Y5 receptors, while NPY3-36 only binds to the Y2 and Y5 receptors[29]. Several studies have shown that activation of the Y1 receptor and injection of NPY can reduce fear memory in rats, while Y2 agonists have no such effect [7, 30, 31]. Moreover, NPY impairs the maintenance of LTP, which is also consistent with our findings [32]. The impairment of memory retrieval and synaptic plasticity caused by inhibiting DPP9 enzyme activity may be related to the activation of Y1 receptors. Y1 receptor activation reduces NMDA-mediated excitatory postsynaptic currents and conversely increases GABAA-mediated inhibitory postsynaptic currents [33].

In addition to participating in biological functions through enzymatic activity, DPP9 also participates in protein-protein interactions [34–36]. We found that DPP9 interacts with the learning- and memory-related proteins Ppp1r9b, Rac1, Ppp2r1a, etc. Rac1 is a small GTPase that strongly affects memory and forgetting [37, 38]. Many studies have shown that actin is an important protein that maintains the morphology and occurrence of dendritic spines and that affecting the aggregation of actin can regulate learning and memory [11, 39–41]. We found that DPP9-interacting proteins are highly involved in aggregation and are localized in the spine, which indicates that DPP9 is likely to be directly involved in the maintenance of direct synaptic connections. By combining data on DPP9-interacting proteins with proteins bidirectionally regulated by DPP9, we found that DPP9 can affect actin aggregation directly or through small GTPases. These findings suggested that DPP9 can affect synaptic viability through multiple pathways and in multiple ways.

Due to the neonatal lethality of DPP9, we were unable to establish an effective knockout mouse strain. Therefore, we used an AAV vector to knock down the target brain region in a more convenient and efficient manner. [42, 43]. CasRx was recently discovered as a knockdown agent. It requires a more specific gRNA, and unlike RNAi, which relies on its own RNA-induced silencing complex, it has lower off-target efficiency and higher knockdown efficiency [44–46].

The hippocampus is generally divided into dorsal and ventral halves. The dorsal side is more involved in learning and memory, while the ventral side is more involved in emotion and emotion regulation. [47]. We modulated DPP9 in the dorsal hippocampus and found that DPP9 did not affect spontaneous or anxious behavior in mice. These findings are consistent with previous reports on the dorsal hippocampus [48, 49]. The strong effect of DPP9 on memory and synaptic viability suggested that DPP9 is critical for memory-related behaviors. Notably, the change in DPP9 did not affect the learning process. These findings indicate that DPP9 regulates the maintenance and extraction of memory and does not participate in the acquisition of acquired information.

We used proteomics to study the mechanism by which DPP9 might be involved in memory and found that DPP9-derived DEPs were highly involved in synaptic function and memory-related pathways, which validated our behavioral and electrophysiological results. Moreover, we also found that there were many differences between the DEPs of DPP9 KD and DPP9 OE. This may be because the effect of OE is better than that of KD. More importantly, we analyzed the DEPs between DPP9 KD and DPP9 OE to determine the enrichment of the pathways separately. These pathways suggest that the DPP9 OE more directly regulates glutamate receptors and synaptic structures.

In a proteomic study on how memantine improved the cognition of triple transgenic mouse Alzheimer’s disease (AD), DPP was found to be the most downregulated protein [50]. Another recent study showed that the downregulation of DPP4 contributes to the improvement of AD symptoms, which is mainly due to the involvement of DPP4 in the cleavage of pyroGlu3–Aβ. [51]. Although DPP4 is not expressed in the brain, the use of DPP4 inhibitors in APP/PS1 mice can also improve AD symptoms. This difference is likely related to changes in the permeability of cerebral blood vessels in AD [52]. Since DPP9 and DPP4 exhibit similar enzymatic activities, DPP9 is likely to play a more important role in the treatment of AD. Furthermore, some studies have confirmed that Aβ activates the NLRP1 inflammasome, thereby triggering severe neuroinflammation and neurotoxicity [53–55]. DPP9 can inhibit NLRP1 through enzymatic activity and FIIND domain binding[56]. Based on these studies, we found that DPP9 seems to play an important role in the treatment of AD, which will also be our future research direction.

This study has several limitations. We did not construct a conditional KO mouse model, which led to our inability to completely knock out DPP9 in the hippocampus. In the future, we will further study how DPP9 affects actin aggregation and small GTPase function.

In summary, we revealed the role and mechanism of DPP9 in fear memory. We found that DPP9 was more highly expressed in hippocampal neurons with high enzyme activity. Its expression increases after fear memory formation. DPP9 knockdown or overexpression attenuated or enhanced fear memory, respectively, by interacting with both NPY and actin regulation proteins. Therefore, hippocampal DPP9 is an important protein that bidirectionally regulates fear memory. These findings will be helpful for the treatment of memory-related diseases in the future.

## Acknowledgments

We would like to thank the Institutional Center for Shared Technologies and Facilities of Kunming Institute of Zoology (KIZ), Chinese Academy of Sciences (CAS), for providing us with mass spectrometry. We are grateful to Lin Zeng for the technical support. We are also grateful to Yu Pan from the Technion - Israel Institute of Technology, who helped us discuss and construct the illustrations. The STI2030-Major Projects (Grant Nos. 2022ZD0204900 to L.X.) The Natural Science Foundation of China (Grant Nos. 32071029 to Q.X.Z., 32271080 to L.X. and 32100835 to X.T.); the Department of Science and Technology Program of Yunnan Province (Grant Nos. 202101AS070052 to Q.X.Z., and 202301AT070298 to J.N.L.)

## Credit author statement

Conceptualization: Y.B. Z and Q.X. Z; Methodology: Y.B. Z; Formal Analysis: Y.B. Z; Investigation: Y.B. Z, S.Z. W, LW; Writing – Original Draft: Y.B. Z; Writing – Review & Editing: Y.B. Z, Q.X. Z and W.T. G; Visualization: Y.B. Z; Supervision: Q.X. Z, L. X and Y.B. Z; Funding Acquisition: L. X and Q.X. Z

---

**Supplemental figure 1.**
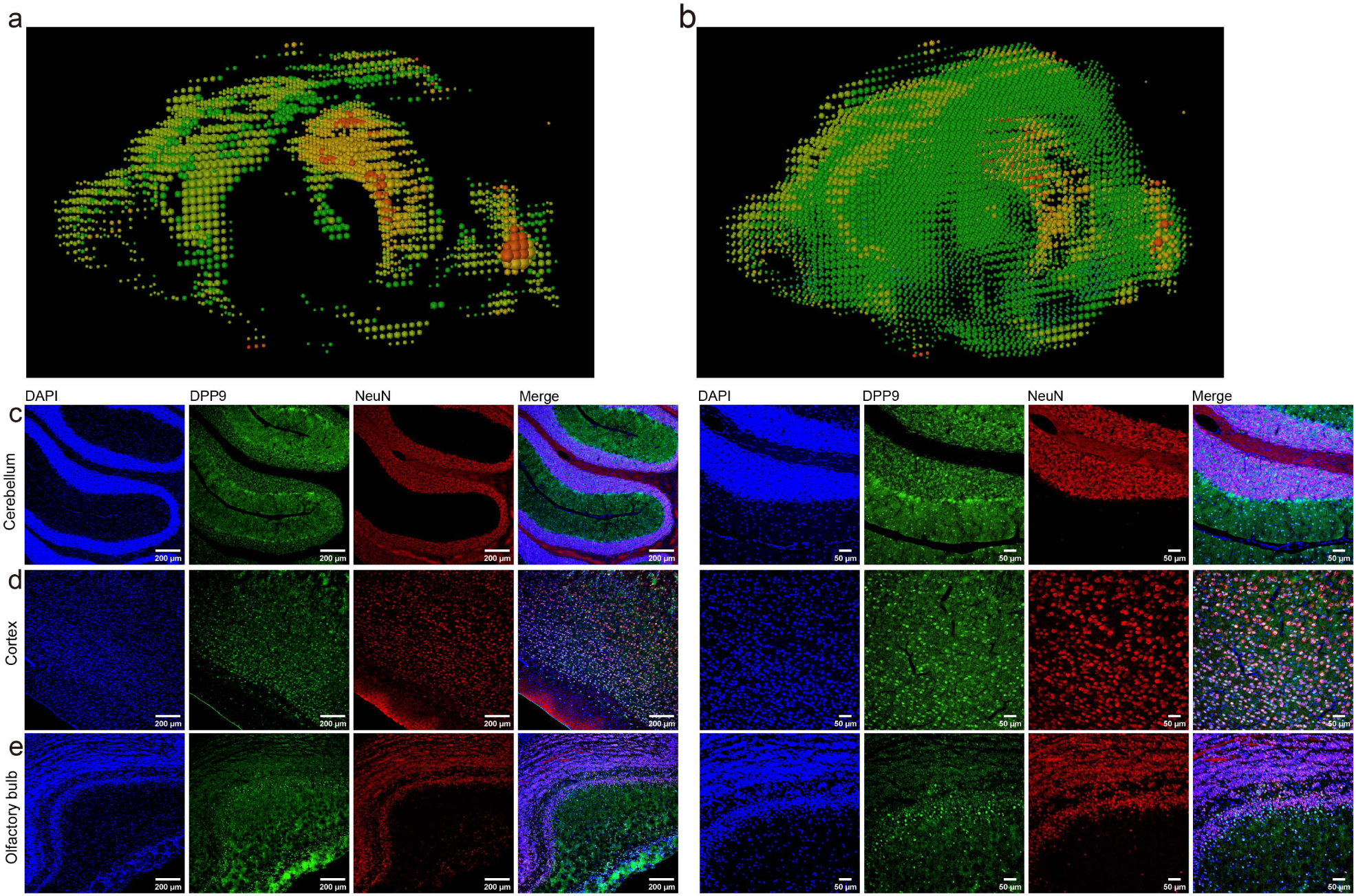
DPP9 mRNA distribution in the brain. (a, b) FISH of the whole-brain distribution of DPP9 mRNA using the Allen Brain Database. (c-d). Immunofluorescence staining of the mouse brain in the cerebellum, cortex, and olfactory bulb. DPP9 (green), NeuN (red), DAPI (blue), 10X (left) Scale 200 μm, 20X (right) Scale 50 μm.

**Supplemental figure 2.**
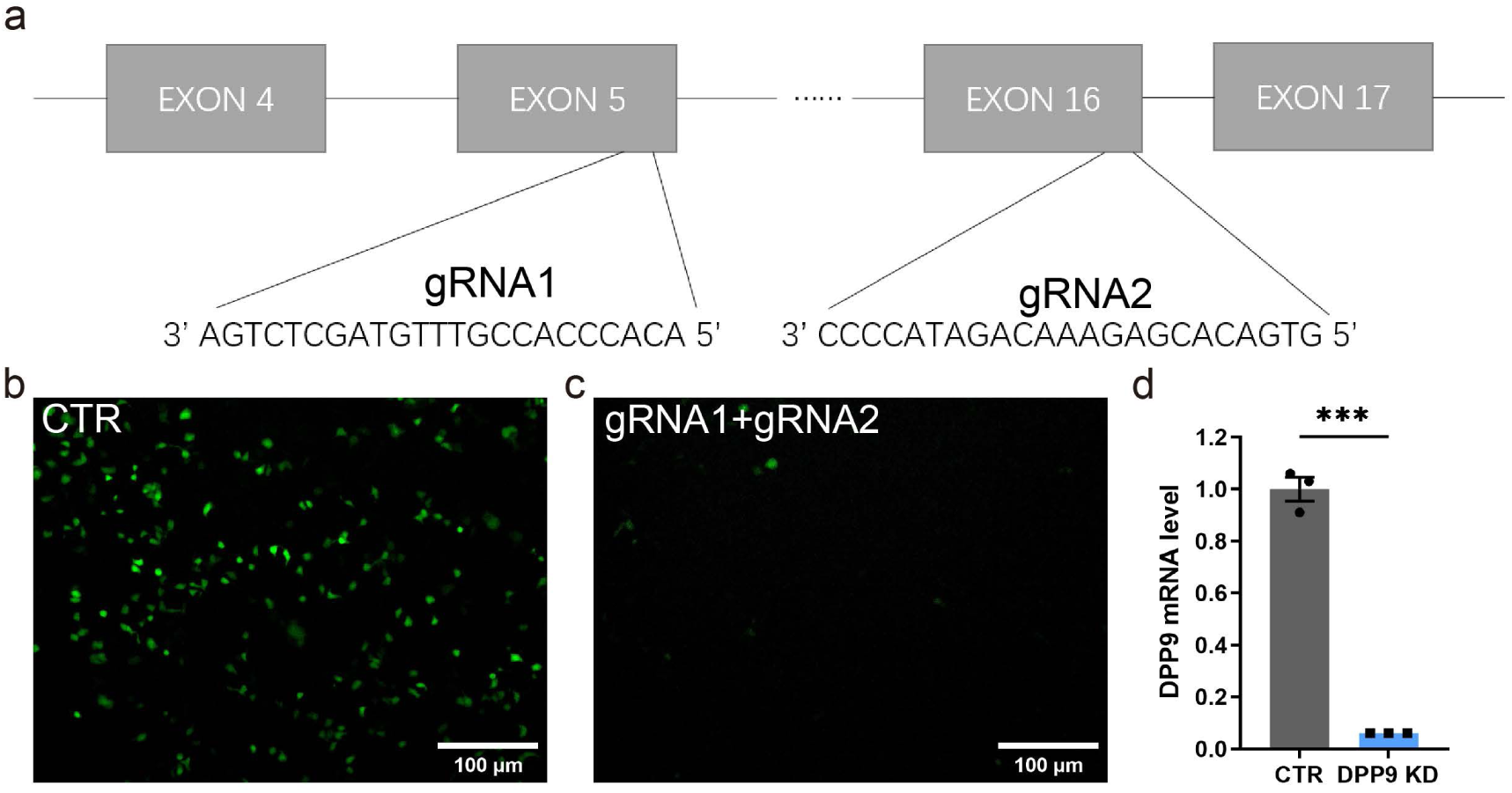
DPP9 CasRX cell verification. (c) Schematic representation of the location of the two gRNAs in the DPP9 genome. (d, e) Fluorescent signal of 293T cells co-transfected with the DPP9 gene (green) and CasRX (left, CRT (random gRNA), right, DPP9 gRNA (gRNA1+gRNA2)). (f) DPP9 mRNA levels after co-transfection of the DPP9 gene and CasRX in 293T cells. t-test, ***P<0.001, mean ± SEM.

**Supplemental figure 3.**
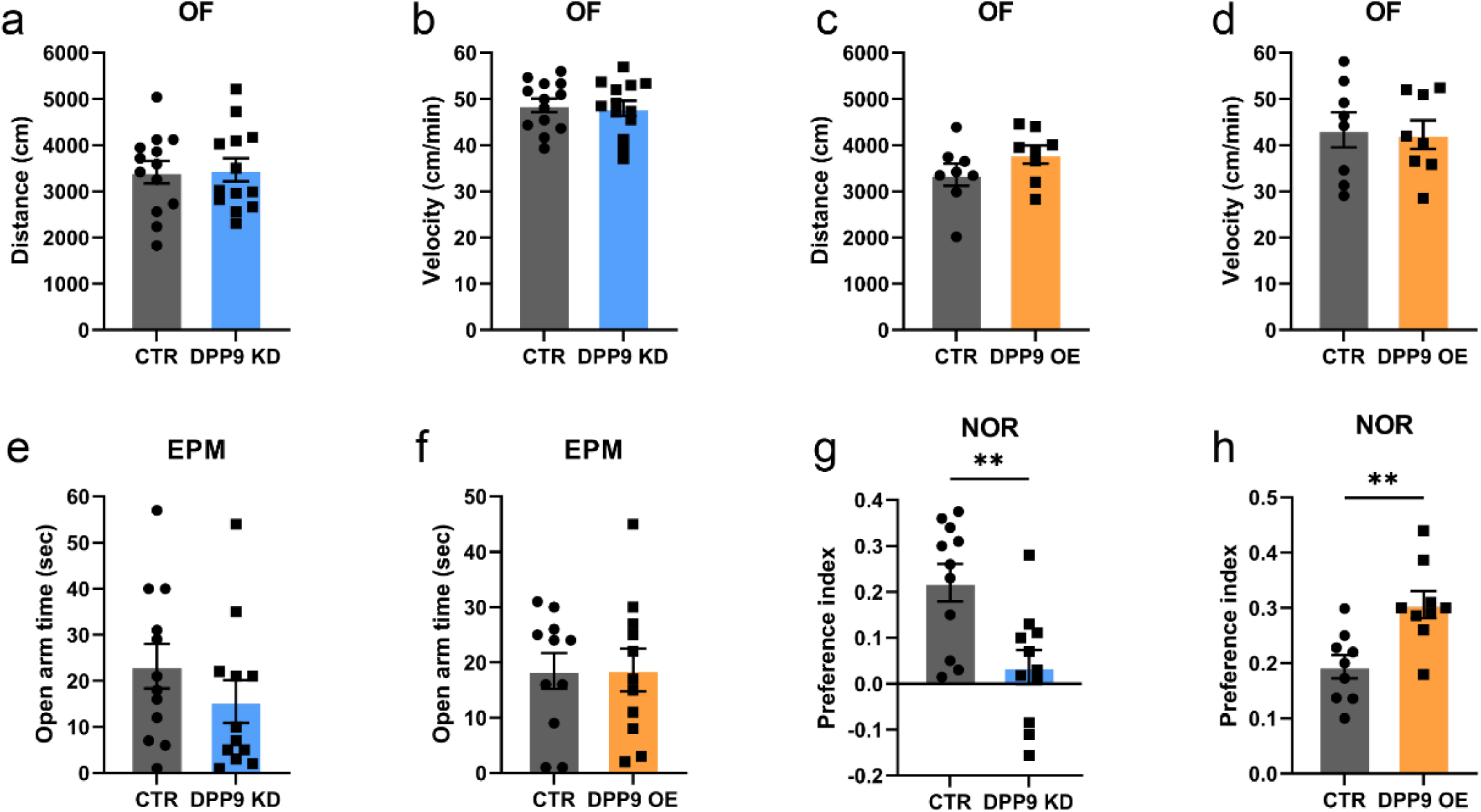
Behavioral experiment on DPP9 knockdown and overexpression. (a, b) Open field test (OF) comparing the distance (left) and velocity (right) for KD and control (CTR). (KD, n=12, OE, n=8, t test) (c, d) OF test comparing the distance (left) and velocity (right) for OE and CTR. (KD, n=12, OE, n=8, t test) (e, f) Elevated plus maze (EPM) test comparing the open arm time for KD and CTR (left), OE and CTR (right). (KD, n=12, OE, n=11, t test) (g, h) Novel object recognition (NOR) test comparing the preference index for KD and CTR (left), OE and CTR (right) (KD, n=11, OE, n=9, t test).

**Supplement figure 4.**
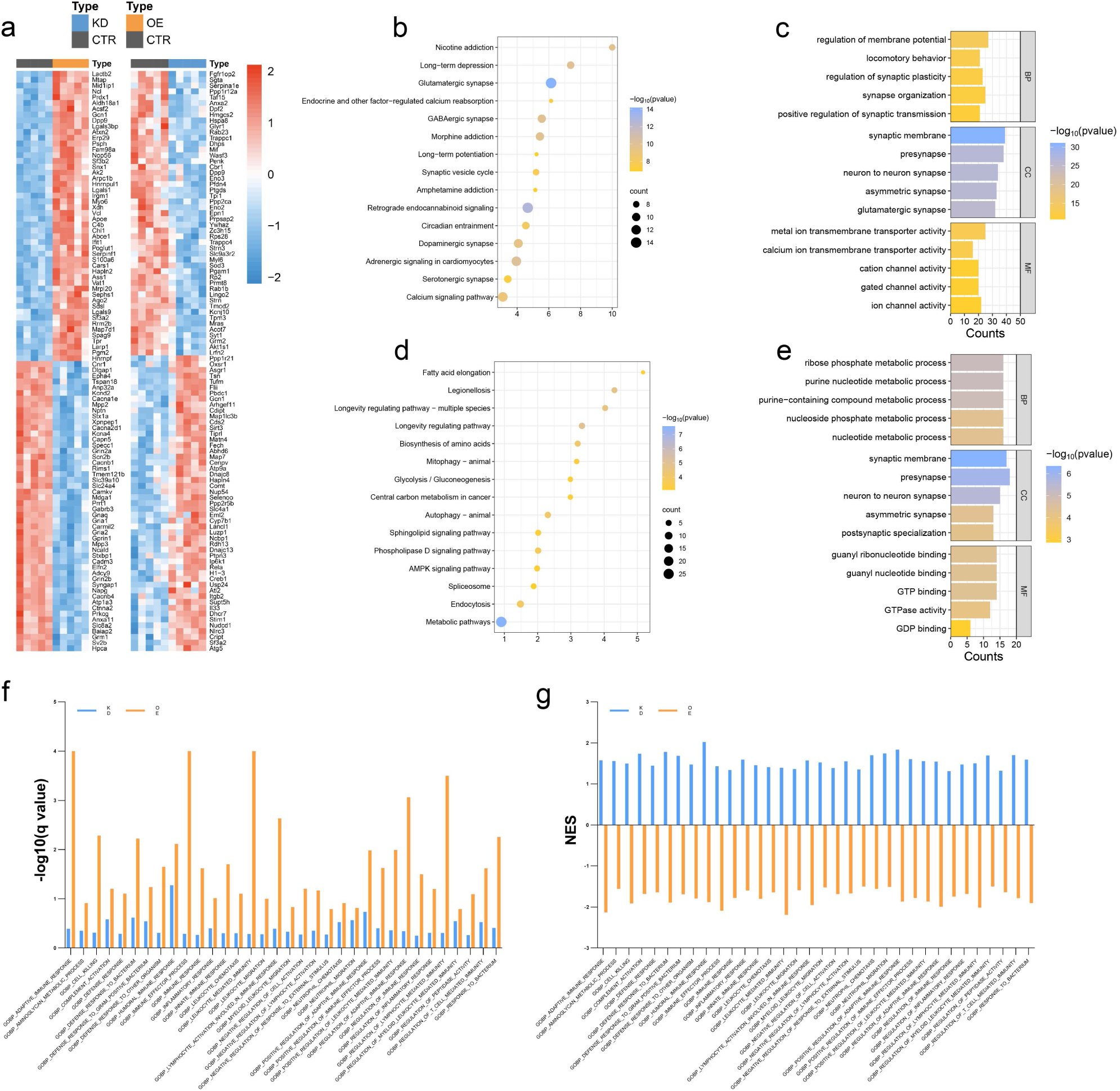
Pathway enrichment of DPP9 OE and KD proteomes. (a) Heatmap of the top 100 proteins with the smallest P value in DPP9 KD and OE. (b, c) KEGG pathway enhancement (left) and GO enhancement (right) for the top 100 proteins with the smallest P value in DPP9 OE. (d, e) KEGG pathway enhancement (left) and GO enhancement (right) for the top 100 proteins with the smallest P value in DPP9 KD. (f, g) GO biological process GSEA of DPP9 KD and OE (left, -log10(q value), right, NES)

**Supplemental figure 5.**
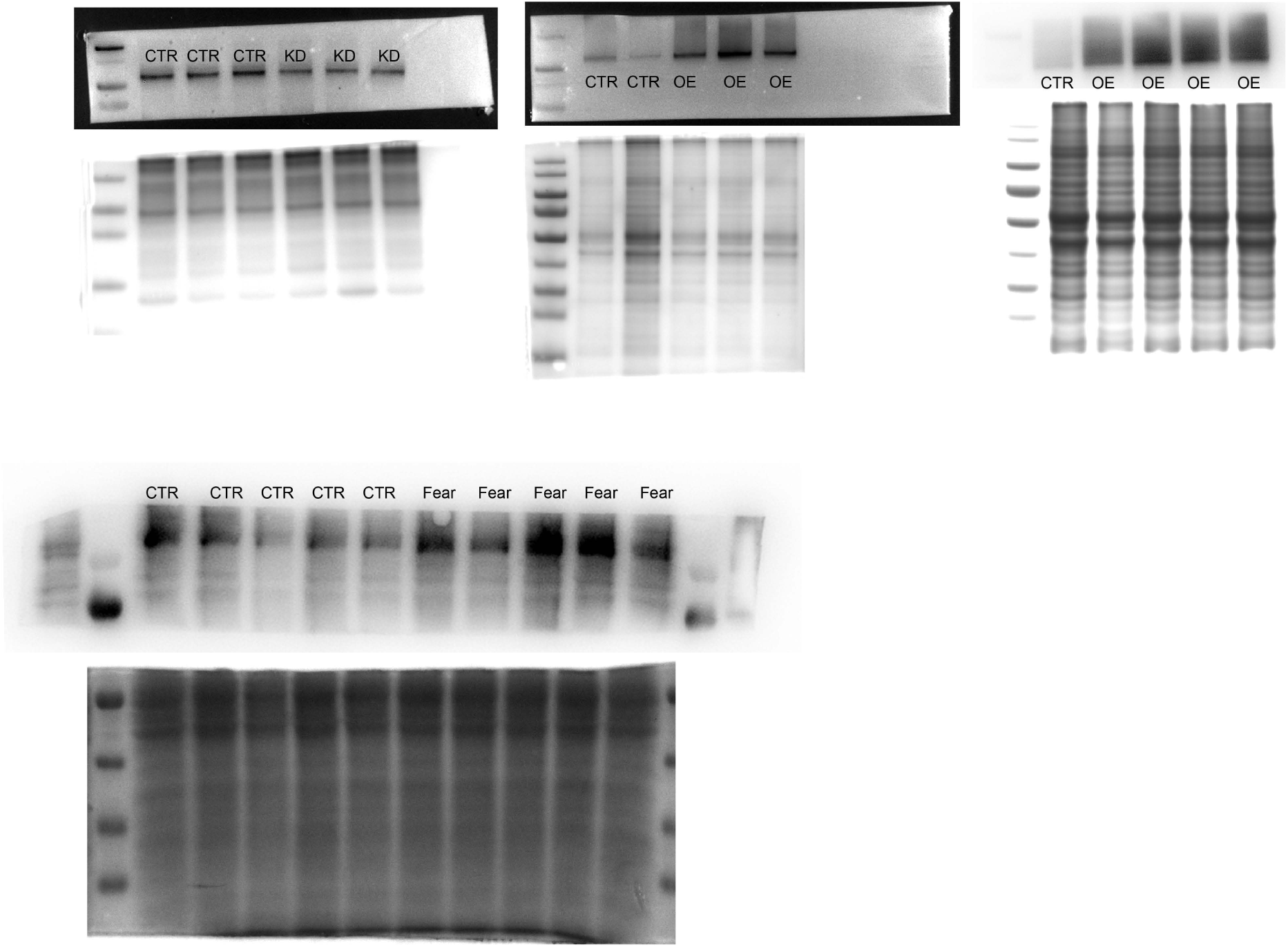
Raw immunoblot image.

